# Investigation on the changes of *Bifidobacterium* and lactic acid bacteria in soybean meal fermented feed

**DOI:** 10.1101/2021.12.16.472915

**Authors:** Lingyu Kang, Huayou Chen, Tao Feng, Keyi Li, Zhong Ni, Ebin Gao, Yangchun Yong

## Abstract

The main objective of this research was to explore the dynamic changes of *Bifidobacterium* and lactic acid bacteria (LAB) in the process of feed fermentation under anaerobic condition, so as to increase the number of fermented bacteria of *Bifidobacterium* from the aspect of strain combination. The results showed that when *Bifidobacterium lactis* (*B. lactis*, i.e. *Bifidobacterium animalis subsp. lactis*) fermented with *Bacillus coagulans* or *Lactobacillus paracasei*, the maximum number of *B. lactis* in those samples was 9.42 times and 4.64 times of that of fermented sample with *B. lactis* only. The soybean meal was fermented by *B. lactis*, *L. paracasei* and *B. coagulans*, and the number of *B. lactis* reached the maximum after fermented 10 days, which was 6.13 times of that in unfermented sample. The reducing sugar content and highest activity of α-galactosidase were higher than the control. These results suggest that *B. coagulans* and *L. paracasei* can promote the growth of *B. lactis*. It is inferred that *B. coagulans* can metabolize normally in aerobic, micro-aerobic and anaerobic environments, consume oxygen, produce digestive enzymes, and cooperate with *L. paracasei* to produce metabolic products benefit for the growth of *B. lactis*.

## 1 Introduction

Fermented feed is rich in small peptides, organic acids and other functional factors. The macromolecule and anti-nutritional factors in raw material are degraded through microbial fermentation, and the quality of fermented feed is directly affected by the evolution of fermentation probiotics. Mixed culture fermentation has become the main fermentation method for the preparation of biological feed. Compare with monoxenie fermentation, mixed culture fermentation can better reflect the synergism and complementarity among microorganisms. Ding et al.(Ding et al., 2020) optimized the fermentation conditions of tea residue, in which fermentation strains were *Bacillus subtilis*, *Aspergillus niger* and *Saccharomyces cerevisiae*, and the crude protein, reducing sugar, and cellulose activity of the fermentation products were significantly improved compared with those before fermentation. Previous studies have indicated that mixed culture fermentation can improve the nutritional quality of feed by reducing antinutritional factors in soybean meal, corn and other raw materials and by increasing nutrient bioavailability(Li et al., 2019; Shi, Zhang, Lu, & Wang, 2017). Mixed culture fermentation system is very complicated with more relationships among fermented probiotics. Chen et al.(Chen, Shih, Chiou, & Yu, 2010) revealed that *Aspergillus oryzae* and *Lactobacillus casei* could degrade the antigenic protein in soybean meal. Nowadays, there are many studies on technology optimization of fermented feed with mixed strains, but there are few reports on the interaction mechanism between fermentation strains. There are abundant metabolites in the several strains coculture environment, and the interaction and metabolic pathways of different strains are different from those of single culture. Therefore, screening the strains with synergistic relationship will be the core content of fermented feed study.

Aerobic fermentation, which is easy to be polluted by pathogenic bacteria in the environment, is charactered with more equipment invest and obvious material loss. Anaerobic fermentation process avoids the waste of raw materials and the possibility of contamination during fermentation. Moreover, anaerobic fermentation process doesn’t need oxygen, so it consumes less energy than aerobic fermentation. Lactic acid bacteria (LAB) are the main fermentation microorganism in anaerobic fermentation process. The growth characteristics and fermentation effect of LAB are quite different with each other. *L. fermentum* and *L. plantarum* are dominating fermentation microorganism(Camu et al., 2007). *L. paracasei* and *L*. rhamnosus have strong ability of acid producing performance and good health function(Demers-Mathieu, St-Gelais, Audy, Laurin, & Fliss, 2016). *B. coagulans* produces both lactic acid and spores, which is favorable to long term storage of feed(Zhou et al., 2020). Compared with *Bacillus subtilis*, *B. coagulans* can metabolize normally in both micro-aerobic and anaerobic environments, consume oxygen and produce lactic acid. Compared with LAB, *B. coagulans* has more abundant enzyme included protease, amylase and xylanase. *Bifidobacterium* is anerobic and can produce a variety of digestive enzymes and produce B vitamins such as folic acid, directly providing nutrients to the animal(Bujna, Farkas, Tran, Dam, & Nguyen, 2018). Meanwhile, *Bifidobacterium* can make good use of stachyose in soybean meal to degrade anti-nutritional factors in raw materials(Grmanová, Rada, Sirotek, & Vlková, 2010), which has been applied in broiler breeding(Li, Lu, Wu, & Lien, 2014). Research suggested that *Bifidobacterium* tend to exhibit weak growth even in milk, and always need long fermentation time and strict condition of anaerobiosis(Abdulamir, Yoke, Nordin, & Abu, 2010). It may be that *Bifidobacterium* are more likely to be affected by factors such as oxygen, pH, storage temperature than other bacteria(Kurmann & Rasic, 1991). *Bifidobacterium* are mostly used as microbial agents added to the diet directly, but studies on *Bifidobacterium* as fermented probiotics for feed are rarely reported.

In this study, soybean meal was used as the main fermentation raw material, and *B. lactis* was mixed with different LAB for fermentation. The microflora changes of different fermentation stages were measured by quantitative PCR (qPCR), and the quality of fermented feed products was detected.

## 2 Materials and methods

### 2.1 Strains and culture methods

*Bacillus subtilis* CGMCC 1.1086, *Saccharomyces cerevisiae* CGMCC 2.1527 and *Lactobacillus rhamnosus* CGMCC 1.577 used in this study were obtained from China General Microbiological Culture Collection Center (CGMCC). *Lactobacillus fermentum*, *Lactobacillus plantarum*, *Lactobacillus paracasei*, *Bacillus coagulans* and *Bifidobacterium animalis subsp. lactis* were preserved strains in our laboratory.

*B. subtilis* was grown in LB medium at 30 °C with a shaking of 180 rpm. *S. cerevisiae* was grown in YPD medium at 28 °C with a shaking of 200 rpm. *Bifidobacterium* and LAB were grown anaerobically in MRS medium at 37°C. When the optical density at 600 nm for the growing cultures was between 0.8 and 1.0, the strains were used in solid-state fermentation.

### 2.2 pH and total acid

Soybean meal 45 g, soybean residue 45 g, and inferior wheatmeal 10 g were used as fermentation medium, and water content was adjusted to 55% after inoculation with a 2% ratio of experimental strains.,. The mixture was then packaged in polythene fermented bags and sealed by heat. All the bags were incubated at 37°C under anaerobic condition. Samples of different fermentation days were taken to test the total acid content and pH value. Total acid content (g of lactic acid*g^-1^) of feed was estimated according to the method Jemaa et al(Ben Jemaa et al., 2017).

### 2.3 Solid-state fermentation process

Soybean meal 45 g, soybean residue 45 g, inferior wheatmeal 10 g were used as fermentation medium. Inoculum amount of *B. subtilis* and *S. cerevisiae* was totally 2%, inoculum rate of *L. fermentum*, *L. plantarum*, *L. paracasei*, *L. rhamnosus* and *B. coagulans* was 2% individually, and water content was adjusted to 55%. The mixture was then packaged in polythene fermented bags and sealed by heat. All the bags were incubated at 37°C under anaerobic condition. Feed samples of different fermentation time were stored at −20°C for further analysis.

Under the above fermentation conditions, *B. lactis* was mixed with different LAB for fermentation. The concrete operation process was shown in Table 1.

**Table 1.** Combination of fermentative strains

| Sample | Fermented strains |
| --- | --- |
| A | *B. subtilis*, *S. cerevisiae*, *B. lactis*, *L. fermentum* |
| B | *B. subtilis*, *S. cerevisiae*, *B. lactis*, *L. plantarum* |
| C | *B. subtilis*, *S. cerevisiae*, *B. lactis*, *L. paracasei* |
| D | *B. subtilis*, *S. cerevisiae*, *B. lactis*, *L. rhamnosus* |
| E | *B. subtilis*, *S. cerevisiae*, *B. lactis*, *B. coagulans* |
| F | *B. subtilis*, *S. cerevisiae*, *B. lactis*, *L. paracasei*, *B. coagulans* |

### 2.4 Primer sets

The BLAST search tool was used to check the specificity of each primer set(Hidaka, Horie, Akao, & Tsuno, 2010; Schwendimann, Kauf, Fieseler, Gantenbein-Demarchi, & Miescher Schwenninger, 2015; Sheu et al., 2010). The primer sequences and amplicon size were shown in Table 2. Primers were synthesized by GENERAL BIOL (China).

**Table 2.**
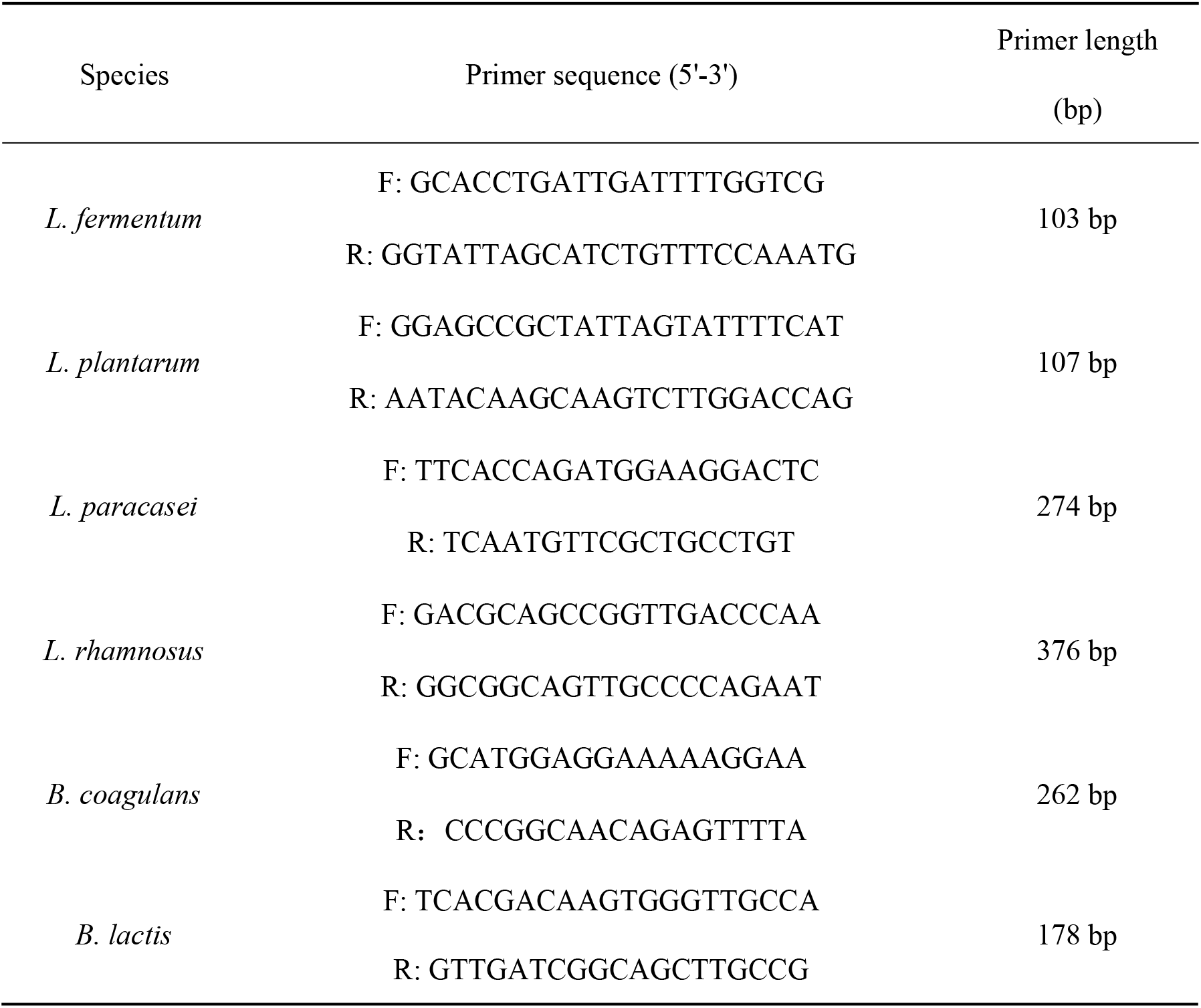
Primer sequence

### 2.5 PCR reaction

The total volume of PCR reaction mixture was 20 μL, with 10 μL Taq Master Mix (Vazyme, China), 2 μL DNA template, 1 μL of each primer and 6 μL ddH_2_O. The PCR programs consisted of the following steps: Denaturation at 95°C for 3 min, followed by 30 cycles of denaturation at 95°C for 30 s, annealing at 60°C for 30 s, and extension at 72°C for 1 min. After completion, extension at 72°C for 10 min. The amplified products were subjected to gel electrophoresis in 3% agarose gels and visualized by ethidium bromide staining.

### 2.6 Preparation of samples

1 g feed sample was put into a 50 mL centrifuge tube with 10 mL PBS buffer and then shaken for 20 min to disperse the microbial cells. After filtration with gauze, 1 mL of the filtrate was centrifuged at 12000 r/min for 2 min, the precipitates were wished with PBS buffer and stored at −20°C. The total DNA of each sample was extracted by FastPure Bacteria DNA Isolation Mini Kit (Vazyme, China).

### 2.7 Quantitative PCR assay

#### 2.7.1 Construction of standard curve

The genomic DNA extracted from standard strains were performed PCR amplification with specific primers. The size of the PCR products was detected by gel electrophoresis in 3% agarose gels, and the bands were accurately cut to obtain the specific fragments of standard strains. The target fragments were purified by Gel Extraction Kit (Sangon Biotech, China). Ten-fold serial dilutions of the purified target fragments, in which concentration was determined, was detected by qPCR. Standard curves were obtained by plotting the Ct values against the target gene copy number. The standard curves of experimental strains were as follows: (1) *L. fermentum*: y=-3.1891x+35.108. (2) *L. plantarum*: y=-3.14x+34.444. (3) *L. paracasei:*y=-3.6016x+37.823. (4) *L. rhamnosus:* y=-3.4245x+35.303. (5) *B. coagulans:*y=-3.4285x+34.501. (6) *B. lactis:* y=-3.6457x+36.8455.

#### 2.7.2 qPCR amplification conditions

qPCR reactions were performed in the QuantStudio 3 Real-Time PCR System (Applied Biosystems, USA). The total volume of qPCR amplification was 20 μL, containing 10 μL of TB Green Premix Ex Taq (Takara, Japan), 6 μL of ddH_2_O, 2 μL of template DNA, 0.8 μL of each primer and 0.4 μL of ROX. Thermal cycling conditions consisted of 1 cycle at 95°C for 30 s, and 40 cycles at 95°C for 5 s followed by 60°C for 30 s. A melting-curve analysis was performed at the end of each qPCR assay

### 2.8 Small peptide

Using Kjeldahl method, refer to GB/T 22492-2008 “Soy peptides power”. The measured small peptide content refers to the proportion of small peptide in the crude protein content of the measured sample. The small peptides consist of oligopeptides, a small number of polypeptides and free amino acids.

### 2.9 α-galactosidase activity

The standard α-galactosidase activity was assayed by the release of ρ-nitrophenol from pNPG. α-galactosidase activity (U/g) of feed was detected according to the method Huang et al(Huang et al., 2018). One unit of enzyme activity (U) was defined as the amount of enzyme releasing 1 μmol of ρ-nitrophenol from10 mmol/L of ρNPG per minute at 37°C with pH 5.5.

### 2.10 Reducing sugar

3,5-Dinitrosalicylic acid (DNS) colorimetric determination of the reducing sugar content in fermented feed was adopted. The details of the analytical process referred to the method proposed by Zhao et al(Zhao, Li, Zhang, Zhang, & Kong, 2013).

### 2.11 Analysis of secondary structure of protein

FTIR absorption spectra from 4000 to 400 cm^-1^ were acquired by Nicolet iS50 (Nicolet, USA). The amide I band performed in the region of 1700-1600 cm^-1^ (Table 3) was analyzed by “PeakFit 4.12” procedure with second-derivative analysis and Fourier self-deconvolution, and “Origin 8.5” procedure was applied to quantify individual component bands according to curve fit. The details of the analytical methods have been reported in previous work(Wang et al., 2014).

**Table 3.** The standard assignment of amide I band

| Secondary structure | Amide I bands/cm<sup>-1</sup> |
| --- | --- |
| β-sheet | 1640~1610 |
| Random coil | 1650~1640 |
| α-helix | 1658~1650 |
| β-turn | 1700~1660 |

### 2.12 Statistical analysis

All results are expressed as mean±standard deviation.

## 3. Results and discussion

### 3.1 Changes of acidity during feed fermentation

As shown in Fig.1(a), the total acid content in feed fermentation showed a gradual growth trend. From 0 to 10 days, the total acid content increased rapidly. As time went on, the total acid content tended to be stable. Among them, *L. rhamnosus* had the strongest acid-forming capacity, and the total acid content up to 5.34%.

**Fig. 1.**
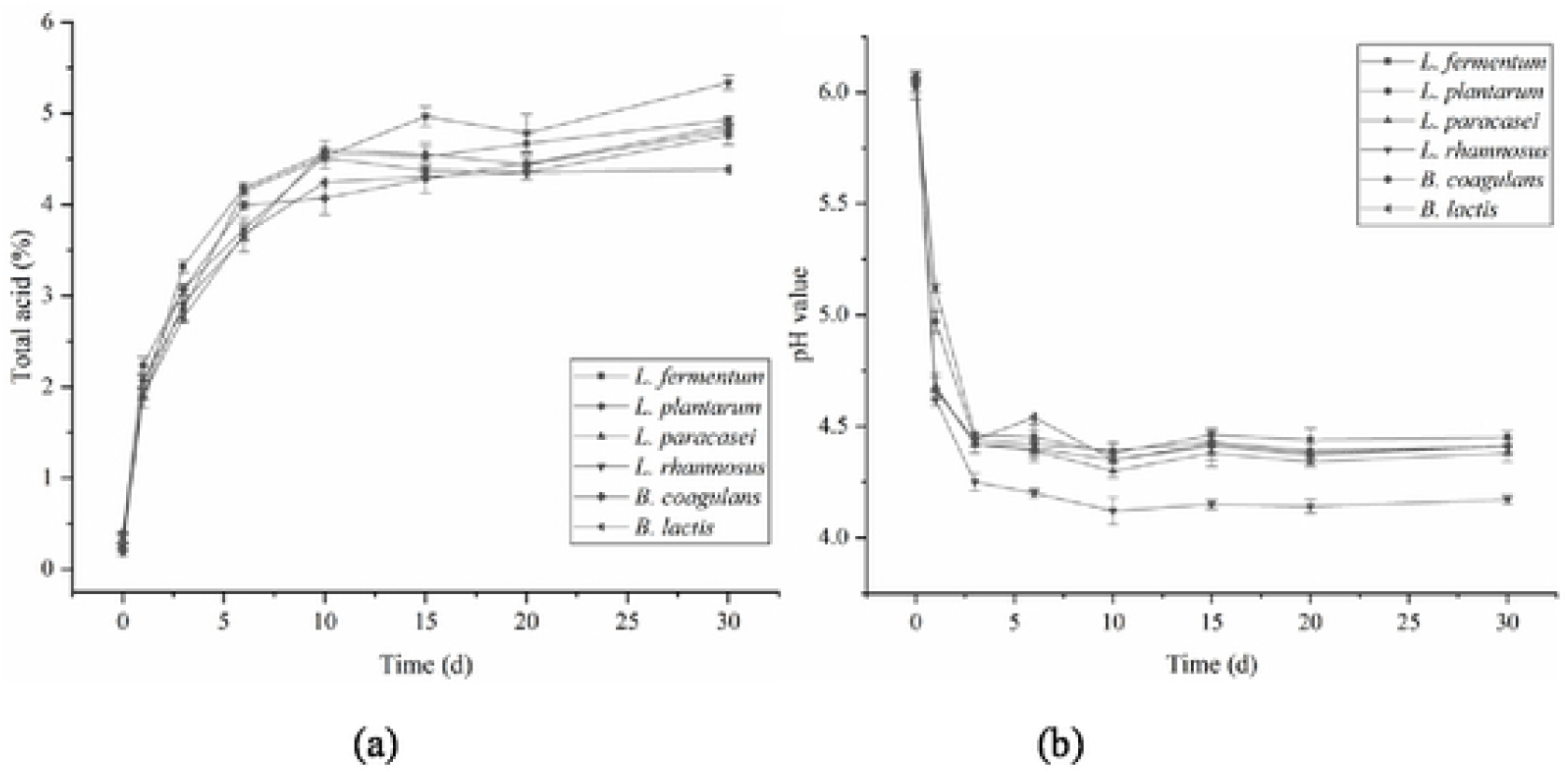
Changes of total acid and pH value in fermentation process.

From Fig.1(b), we can easily see that due to the blooming of LAB, and organic acid produced constantly, the pH value of fermentation products has decreased since the start of fermentation. Over time, the pH value of 6 group all decreased, but the reduced degree was different. The feed inoculated with *L. rhamnosus* had a stable pH ranged from 4.0 to 4.2 after fermentation for 6 days. The pH value of other groups was ranged from 4.3 to 4.5 after fermentation for 30 days, which was basically consistent with the change of total acid in fermentation process.

### 3.2 Changes of *B. lactis* and LAB during feed fermentation

As shown in Fig.2(a), the biomass of *L. fermentum* and *L. plantarum* both reached their maximum on the third day, for 1.84×10^7^ copies/g and 8.88×10^7^ copies/g, respectively. Along with the accumulation of organic acid, pH in the feed declined, and the growth of the bacteria itself was inhibited, but the quantity of bacteria decreased slowly in the fermentation anaphase. The biomass of *L. paracasei* and *L. rhamnosus* reached their maximum on the first day, for 1.84×10^7^ copies/g and 8.88×10^7^ copies/g, respectively. *B. coagulans* also reached its maximum value (1.27×10^7^ copies/g) on the first day, and was 3.74 times as much as those before fermentation. In the early time of fermentation, *B. coagulans* mostly existed in the form of vegetative cell, but the spores were gradually generated as the time went on, leading to the quantity decrease of *B. coagulans*. Although *B. coagulans* could form spores with strong stress resistance, its growth was still affected to some extent. *B. lactis* grew slowly in the early time of fermentation, and reached the maximum value (5.54×10^6^ copies/g) on the tenth day, which was 3.15 times as much as that before fermentation, and then decreased gradually and maintained at a certain level.

**Fig.2.**
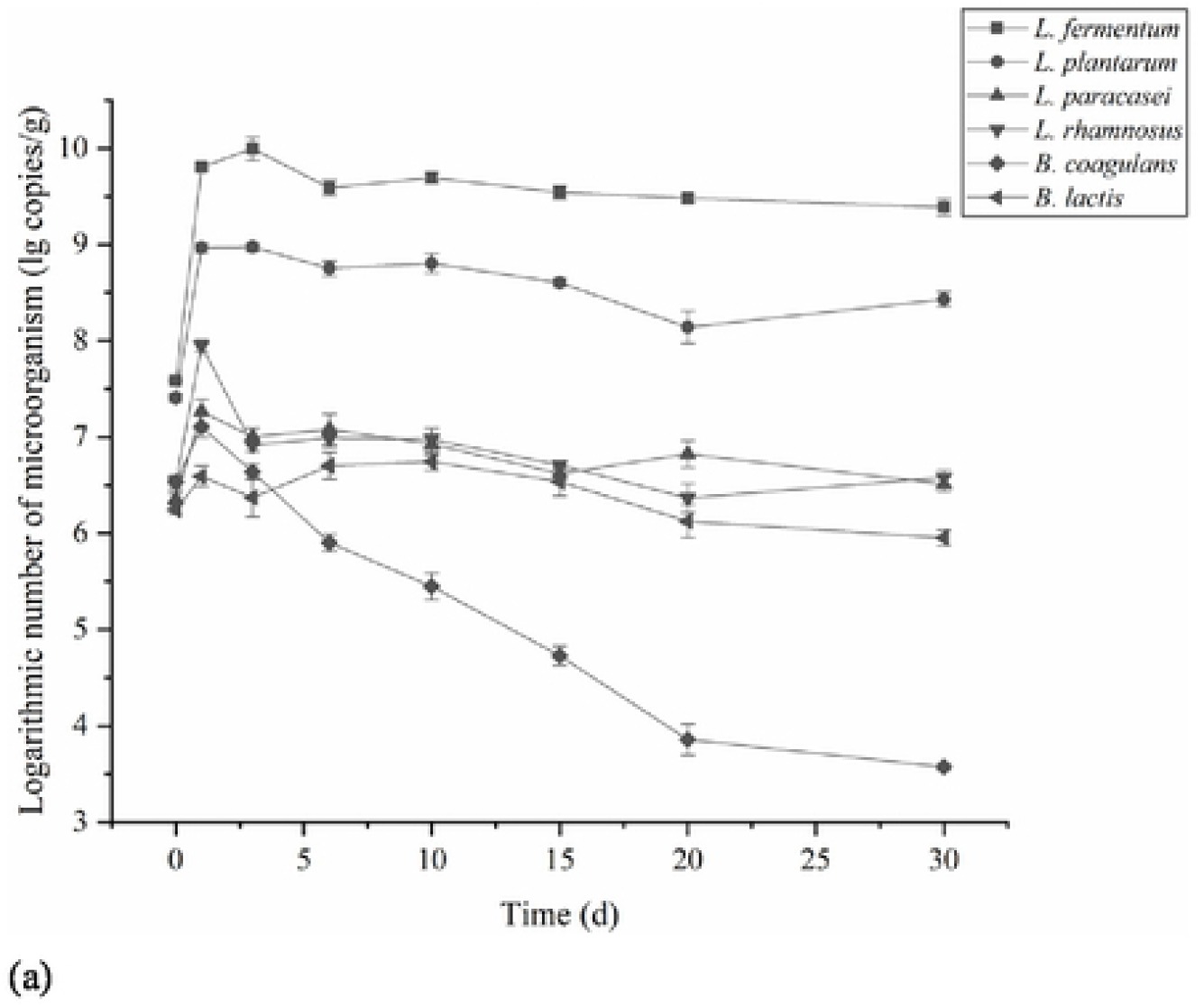
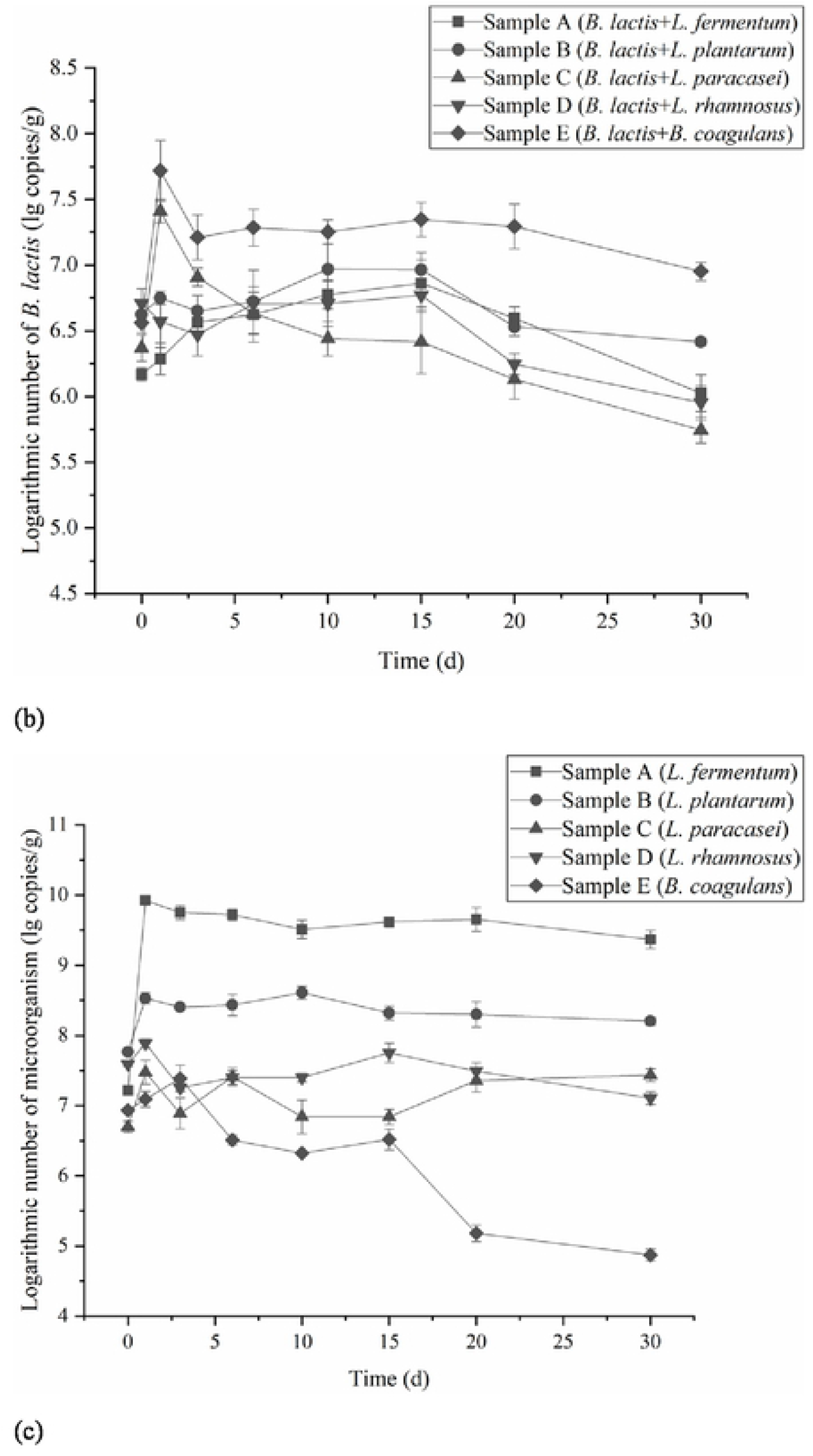
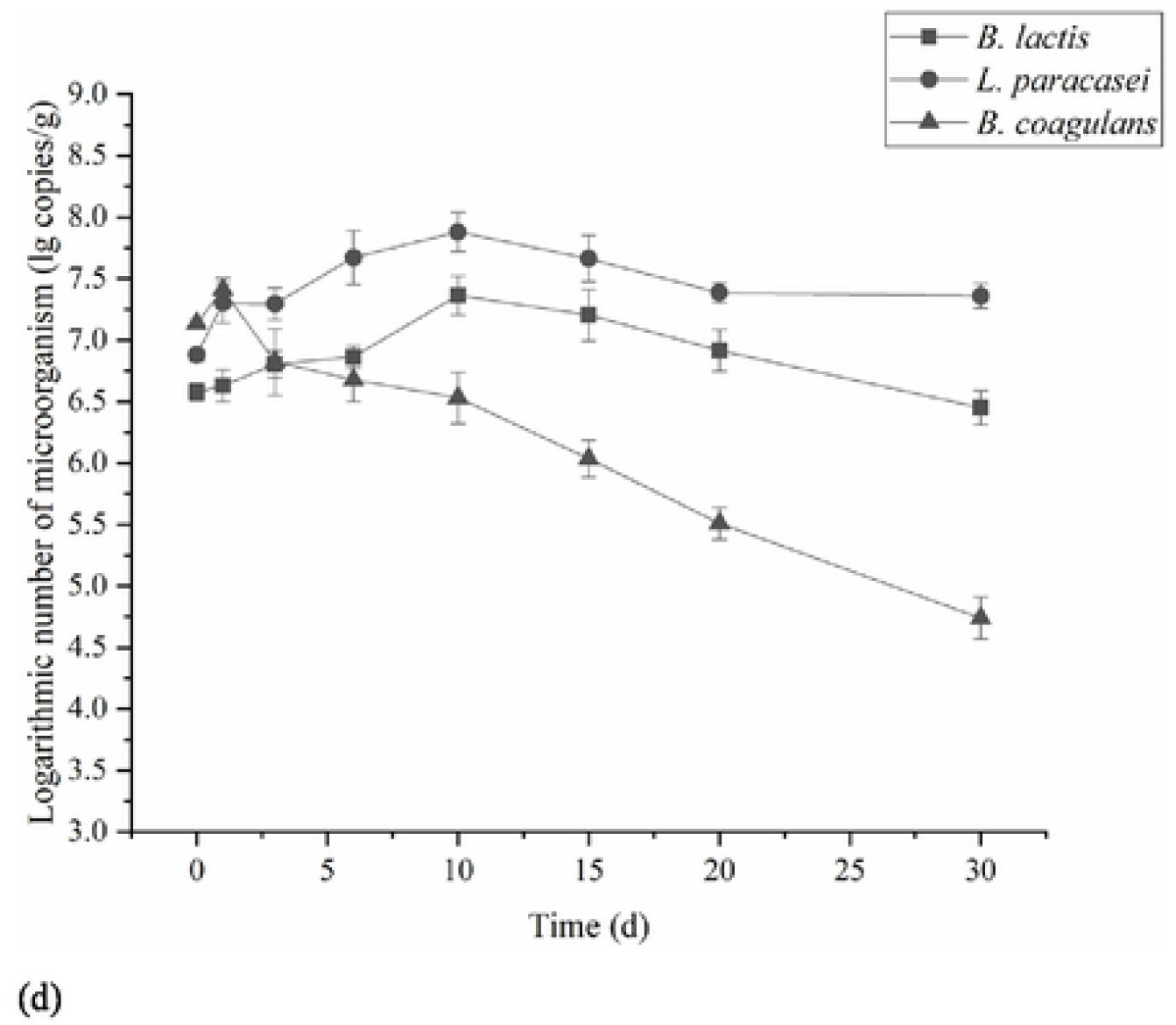
Changes of microflora in fermentation process:(a) fermentation with 6 strain in this study respectively based on *B. subtilis* and *S. cerevisiae*; (b) and (c): fermentation with *B. lactis* and different LAB based on A *sub lilts* and *S. cerevisiae*; (d) sample F: fermentation with *B. lactis*, *L. paracasei* and *B. coagulans* based on *B. subtilis* and *S. cerevisiae*.

Changes of microflora in fermented feed with *B. lactis* and each LAB individually were shown in Fig.2(b) and Fig.2(c). We can easily find that *B. lactis* in sample A reached the maximum value (7.26×10^6^ copies/g) on the fifteenth day of fermentation, which was 4.94 times of that before fermentation, indicating that *L. fermentum* had no inhibitory effect on *B. lactis. B. lactis* reached the maximum on the tenth day of fermentation (9.35×10^6^ copies/g), which was 2.21 times of the number of bacteria before fermentation. *L. plantarum* played a dominant role in fermentation, and grew rapidly as the early stage, and used the nutrients in raw materials preferentially. Interspecific competition may lead to the growth restriction of *B. lactis*. In sample D, *B. lactis* grew slowly and the peak value of bacteria quantity was 1.16 times the number of that before fermentation. *L. rhamnosus* is rich in acid, which resulted in lower pH of fermentation medium. Environmental pH was an important factor affecting the growth of *Bifidobacterium*, the decomposition rate of carbohydrate depends on 6-phosphofructokinase under anaerobic conditions, and the enzyme activity influences glycolytic and lactic acid flux(Li et al., 2016). When *Bifidobacterium* grows in meta-acidic environment, 6-phosphofructokinase is feedback inhibited or repressed(Desjardins, Roy, & Goulet, 1990), so the activity of *Bifidobacterium* is affected, which is not conducive to its growth. Both samples C and E reached their peak values on the first day of fermentation, and their bacterial biomass was 10.98 and 14.30 times of that before fermentation, respectively, which accelerated the fermentation process. *L. paracasei* and *B. coagulans* could promote the growth of *B. lactis* to some extent.

From Fig.2(c), we see that, under anaerobic fermentation conditions, *L. fermentum*, *L. plantarum*, *L. paracasei*, *L. rhamnosus* and *B. coagulans* were inoculated with *B. lactis* individually. The number of *L. fermentum* was 1.66×10^7^ copies/g before fermentation (0 d), and reached its peak value (8.37×10^9^ copies/g) on the first day. The number of *L. plantarum* before fermentation was 5.84×10^7^ copies/g, and reached the maximum on the tenth day (4.06×10^8^ copies/g). *L. paracasei* and *L. rhamnosus* both reached the maximum value on the first day with 2.98×10^7^ copies/g and 7.71×10^7^ copies/g, respectively. *B. coagulans* reached the peak value (1.23×10^7^ copies/g) on the third day of fermentation. After 15 days of fermentation, the number of *B. coagulans* decreased significantly due to the decrease of vegetative cells and spore formation.

In the mixed-culture fermentation process of *B. lactis*, *L. paracasei* and *B. coagulans*, the changes of bacterial biomass are shown in Fig.2(d). *B. lactis* reached the maximum value on the tenth day of fermentation with 2.31×10^7^ copies/g, which was 6.13 times of that before fermentation, and increased compared with that of single fermentation. *L. paracasei* also reached maximum (7.58×10^7^ copies/g) on tenth day and then maintained at a certain level. *B. coagulans* reached the maximum value (2.59×10^7^ copies/g) on the first day of fermentation, and then the bacteria quantity gradually decreased. Oxygen and acidity are the key factors affecting the survival of *Bifidobacterium*. *B. coagulans* not only has the characteristics of *Bacillus* but also produces lactic acid. *B. coagulans* consumed part of oxygen and yielded protease during the early time of fermentation, which provided a good growth environment for *Bifidobacterium*. The growth of *L. paracasei* was stable, and the acid production was suitable. It seemed that *L. paracasei* had no obvious competitive inhibition effect on *B. lactis* when pH of feed environment was reduced. The results showed that the synergistic fermentation of *B. lactis*, *L. paracasei* and *B. coagulans* had application value.

### 3.3 Changes of small peptide, α-galactosidase activity and reducing sugar content during feed fermentation

The changes of small peptide contents in mixed fermentation process of *B. lactis*, *L. paracasei* and *B. coagulans* was shown in Fig.3. The content of small peptides increased rapidly after 3 days of fermentation. It is possible that strains were in exponential phase at initial fermentation, with strong activity and rapid growth. At the same time, protease was produced to hydrolyze the protein in the feed. The content of small peptides increased slowly after 15 days of fermentation and reached 22.65% after 30 days of fermentation.

**Fig.3.**
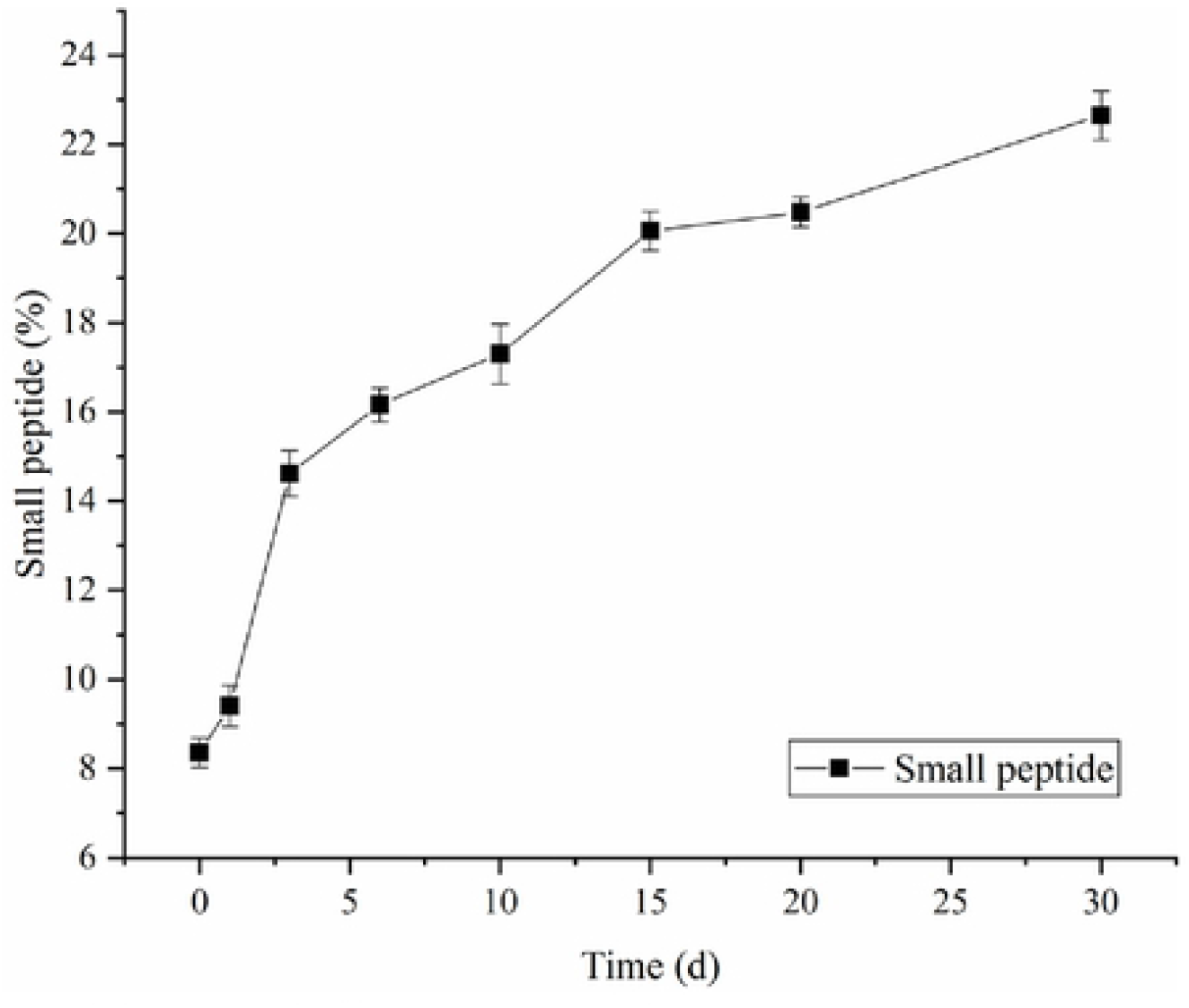
Changes of small peptides during fermentation.

*Bifidobacterium* and *Lactobacillus* species can produce different levels of α-galactosidase and hydrolyze various glycosidic anti-nutritional factors. Studying on the activity of α-galactosidase in the fermentation process can make the correct planning and design of fermentation process. As shown in Fig.4, strains were in exponential phase at initial fermentation, and the enzyme activity gradually increased. The highest enzyme activity of control and sample F were 4.608 U/g and 7.378 U/g respectively after fermentation. In addition, the control maintained relatively stable enzyme activity during stationary phase, and α-galactosidase activity of sample F decreased and was lower than that of control during fermentation anaphase. It is possible that α-galactosidase in various sources had different properties such as protease resistance and galactose tolerance. With the accumulation of metabolites, α-galactosidase inhibitors in the fermentation substrate might influenced the enzyme activity(Huang et al., 2018).

**Fig.4.**
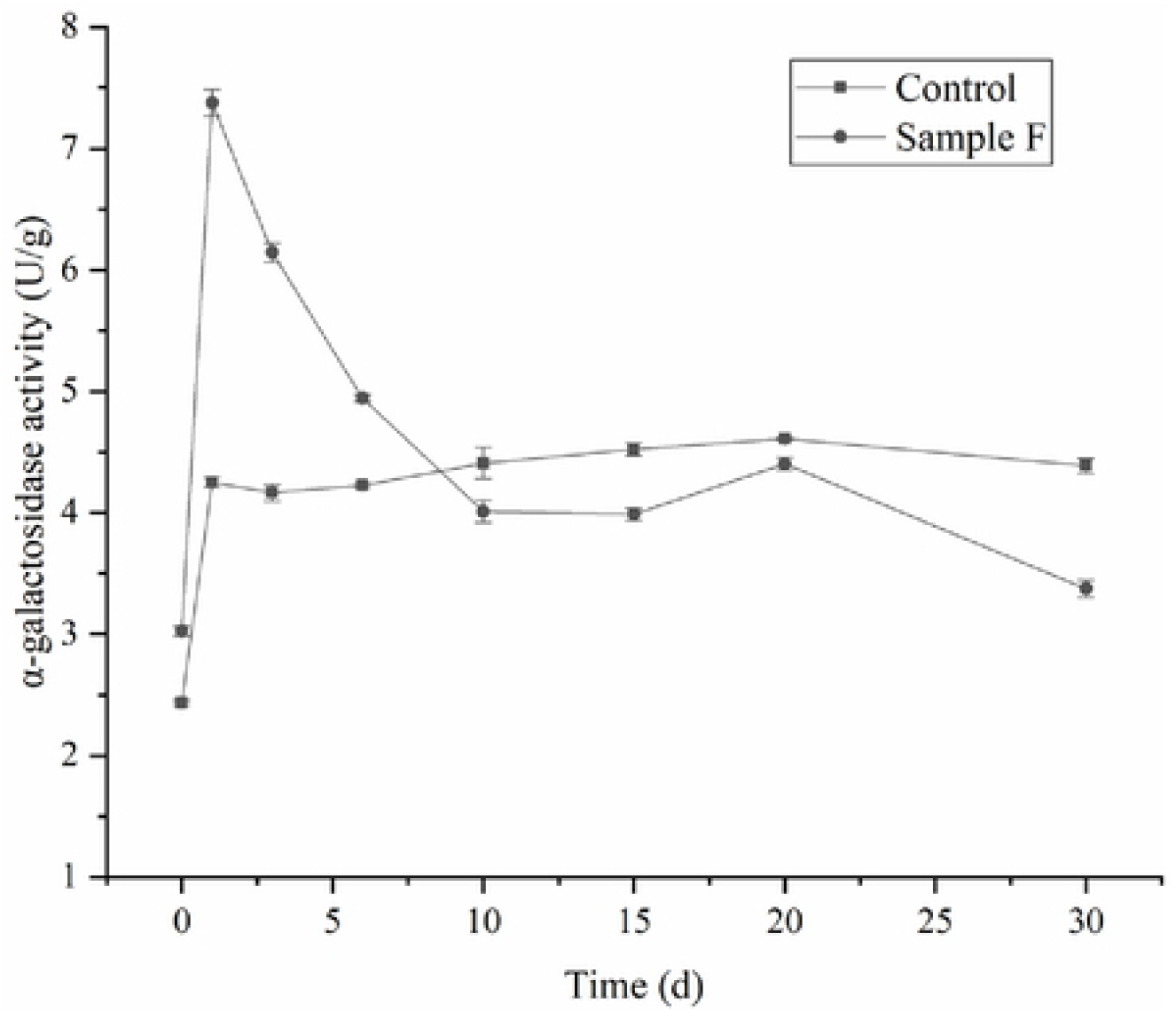
Changes of α-galaclosidasc activity during fermentation: **Control** fermentation with *B. subtilis*, *S. cerevisiae* and *B. lactis*; **Sample F** fermentation with *B. subtilis*, *S. cerevisiae*, *B. lactis*, *L. paracasei* and *B. coagulans*

The changes of reducing sugar content during fermentation were shown in Fig.5. Trend of reducing sugar in sample F was consistent with the control, and fermentation microorganisms transformed the saccharides in feed raw materials into alcohols, organic acids and other substances, and the reducing sugar content gradually decreased as fermentation process went on. From 0 to 6 days, the microorganism grew and multiplied rapidly, and consumed a lot of reducing sugar. After fermentation for 15 days, the reducing sugar content of control and sample F increased to 6.56 mg/g and 8.44 mg/g respectively, and held steady gradually. After fermentation for 30 days, the reducing sugar contents of control and sample F were 7.14 mg/g and 8.00 mg/g, respectively. Microorganism yielding glycosidase and amylase in the fermentation process to hydrolyze non-reducing sugar in feed into glucose and other little molecular carbohydrates to provide energy for the growth of microorganisms. The reducing sugar content of sample F was generally higher than that of control. Sample F had high α-galactosidase activity, which degraded the non-reducing sugar such as raffinose and stachyose into reducing sugars. Meanwhile, as the number of microorganisms gradually stabilized, enzyme activity decreased and reducing sugar consumption also decreased.

**Fig.5.**
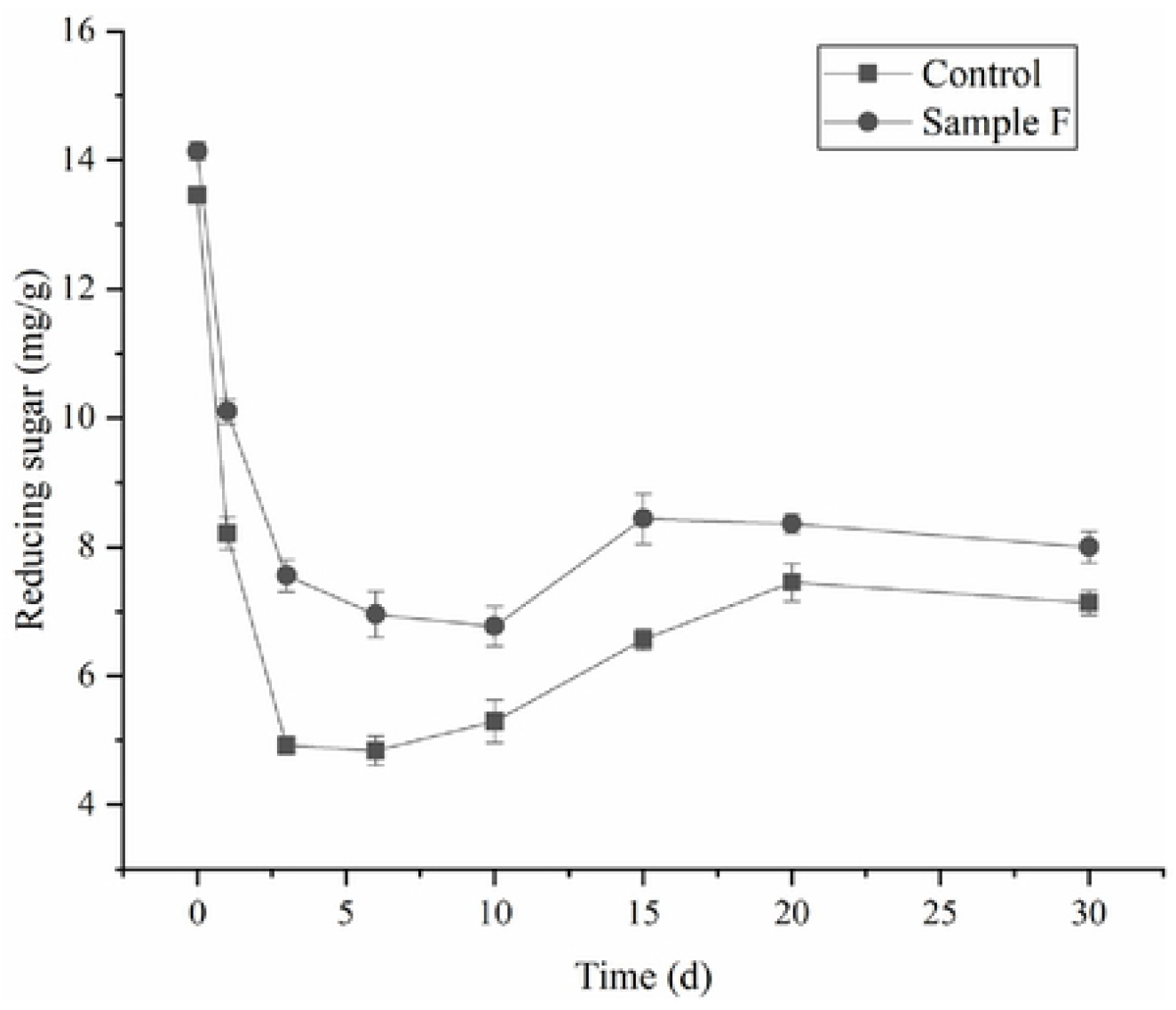
Changes of reducing sugar content during fermentation: **Control** fermentation with *B. subtilis*, *S. cerevisiae* and *B. lactis*; **Sample F** fermentation with *B. subtilis*, *S. cerevisiae*, *B. lactis*, *L. paracasei* and *B. coagulans*

### 3.4 Changes of distinct secondary structure elements of protein in fermented feed

The decomposition of the amide I band was performed in the region of 1700-1600cm^-1^, and the amide I vibration arises mainly from the C=O stretching vibration of protein backbone(Barth, 2007). Amide I vibration is sensitive to secondary structure. A second-derivative analysis and Fourier self-deconvolution were used to resolve and quantify secondary structural components of feed protein. As shown in Table 4, The proportion of secondary structure of protein in the microbial fermented feed (Sample F) was different from that of the unfermented feed. The β-sheet content of the unfermented soybean meal was the highest, which was 34.53%, followed by β-turn, which was 30.08%, and the content of α-helix and random coil was lower, which were 16.30% and 19.10%, respectively, α-helix / β-fold was 47.21%. The ratio of α-helix in fermented feed decreased by 5.07%, and the ratio of β-turn increased by 5.15%. There was little difference in β-sheet and random coil, and the ratio of α-helix / β-folding decreased. It was reported that the decrease of α-helix component may be related to the increase of protein digestibility(Wang et al., 2014). New peak appeared after fermentation (Fig.6), which was speculated to be due to the increase of small peptide content during feed fermentation, leading to protein recombination(Yasar, Tosun, & Sonmez, 2020).

**Fig.6.**
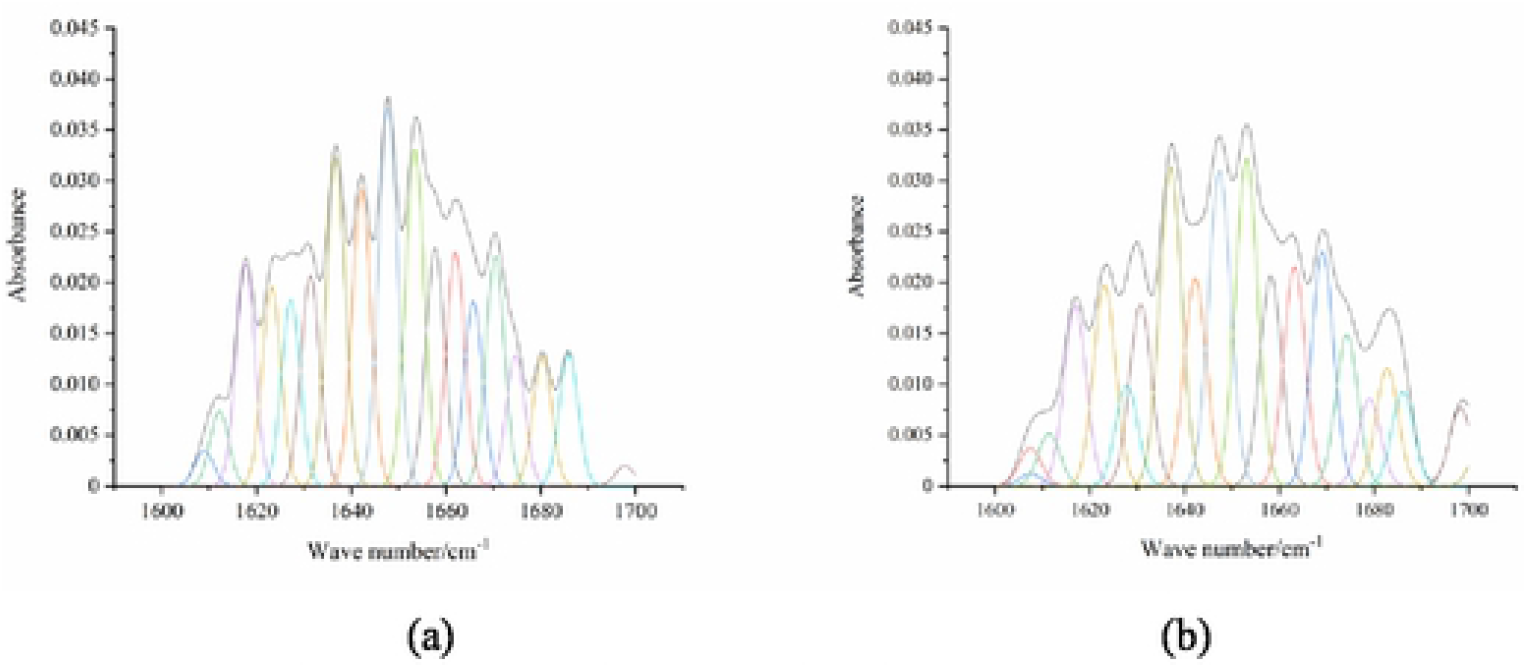
The amide I band curve-fitting results of protcins:(a) fermentation raw material; (b) fermented feed

**Table 4.** Changes of protein secondary structure relative content

| Secondary structure (%) | Before fermentation | After fermentation |
| --- | --- | --- |
| α-helix | 16.30±1.31 | 11.23±1.18 |
| β-sheet | 34.53±1.43 | 35.60±0.92 |
| β-turn | 30.08±1.15 | 35.23±1.78 |
| random coil | 19.10±0.87 | 17.94±1.04 |
| α- helix/β- sheet | 47.21±0.02 | 31.54±0.04 |

## 4 Conclusion

In this study, *L. rhamnosus* had the strongest acid producing ability. When *B. lactis*, *L. paracasei* and *B. coagulans* were used to ferment soybean meal feed together, the peak biomass of *B. lactis* was 6.13 times of that before fermentation, which was higher than sample fermented with *B. lactis* only (3.15 times). The peak value of α-galactosidase activity was 1.6 times higher than the control. On the thirtieth day of fermentation, the content of reducing sugar was 8.00 mg/g, and the content of small peptides was 22.65%, which was 2.7 times of that before fermentation, the proportion of secondary structure of protein in feed also changed. These results suggested that *L. paracasei* and *B. coagulans* could promot the growth of *B. lactis*, and fully ferment soybean meal feed with a synergistic effect.

## Acknowledgements

This study was financially supported by the Open Funding Project of the State Key Laboratory of Biochemical Engineering, China (2018KF-02), Key Research and Development Program (Social Development) of Zhenjiang City (SH2020021).

## REFERENCES

Abdulamir, A. S., Yoke, T. S., Nordin, N., & Abu, B. F. (2010). Detection and quantification of probiotic bacteria using optimized DNA extraction, traditional and real-time PCR methods in complex microbial communities. African Journal of Biotechnology, 9(10), 1481–1492. http://doi.org/10.5897/AJB09.1322.

Barth, A. (2007). Infrared spectroscopy of proteins. Biochimica et Biophysica Acta (BBA) - Bioenergetics, 1767(9), 1073–1101. http://doi.org/ https://doi.org/10.1016/j.bbabio.2007.06.004.

Ben Jemaa, M., Falleh, H., Neves, M. A., Isoda, H., Nakajima, M., & Ksouri, R. (2017). Quality preservation of deliberately contaminated milk using thyme free and nanoemulsified essential oils. Food Chemistry, 217, 726–734. http://doi.org/ https://doi.org/10.1016/j.foodchem.2016.09.030.

Bujna, E., Farkas, N. A., Tran, A. M., Dam, M. S., & Nguyen, Q. D. (2018). Lactic acid fermentation of apricot juice by mono- and mixed cultures of probiotic Lactobacillus and Bifidobacterium strains. Food Science and Biotechnology, 27(2), 547–554. http://doi.org/10.1007/s10068-017-0269-x.

Camu, N., De Winter, T., Verbrugghe, K., Cleenwerck, I., Vandamme, P., Takrama, J. S.,… De Vuyst, L. (2007). Dynamics and Biodiversity of Populations of Lactic Acid Bacteria andAcetic Acid Bacteria Involved in Spontaneous Heap Fermentation of Cocoa Beans inGhana. Applied and Environmental Microbiology, 73(6), 1809–1824. http://doi.org/10.1128/AEM.02189-06.

Chen, C. C., Shih, Y. C., Chiou, P. W. S., & Yu, B. (2010). Evaluating Nutritional Quality of Single Stage- and Two Stage-fermented Soybean Meal. Asian-Australasian Journal of Animal Sciences, 23(5), 598–606.

Demers-Mathieu, V., St-Gelais, D., Audy, J., Laurin, É., & Fliss, I. (2016). Effect of the low-fat Cheddar cheese manufacturing process on the viability of Bifidobacterium animalis subsp. lactis, Lactobacillus rhamnosus, Lactobacillus paracasei/casei, and Lactobacillus plantarum isolates. Journal of Functional Foods, 24, 327–337. http://doi.org/10.1016/j.jff.2016.04.025.

Desjardins, M., Roy, D., & Goulet, J. (1990). Growth of Bifidobacteria and Their Enzyme Profiles. Journal of Dairy Science, 73(2), 299–307. http://doi.org/ https://doi.org/10.3168/jds.S0022-0302(90)78673-0.

Ding, X., Yao, L., Hou, Y., Hou, Y., Wang, G., Fan, J., & Qian, L. (2020). Optimization of Culture Conditions During the Solid-State Fermentation of Tea Residue Using Mixed Strains. Waste and Biomass Valorization, 11(12), 6667–6675. http://doi.org/10.1007/s12649-019-00930-4.

Grmanová, M., Rada, V., Sirotek, K., & Vlková, E. (2010). Naturally occurring prebiotic oligosaccharides in poultry feed mixtures. Folia Microbiologica, 55(4), 326–328. http://doi.org/10.1007/s12223-010-0050-5.

Hidaka, T., Horie, T., Akao, S., & Tsuno, H. (2010). Kinetic model of thermophilic l-lactate fermentation by Bacillus coagulans combined with real-time PCR quantification. Water Research, 44(8), 2554–2562. http://doi.org/10.1016/j.watres.2010.01.007.

Huang, Y., Zhang, H., Ben, P., Duan, Y., Lu, M., Li, Z., & Cui, Z. (2018). Characterization of a novel GH36 α-galactosidase from Bacillus megaterium and its application in degradation of raffinose family oligosaccharides. International Journal of Biological Macromolecules, 108, 98–104. http://doi.org/ https://doi.org/10.1016/j.ijbiomac.2017.11.154.

Kurmann, J. A., & Rasic, J. L. (1991). The Health Potential of Products Containing Bifidobacteria. Therapeutic Properties of Fermented Milks, 117–158.

Li, C., Lu, J., Wu, C., & Lien, T. (2014). Effects of probiotics and bremelain fermented soybean meal replacing fish meal on growth performance, nutrient retention and carcass traits of broilers. Livestock Science, 163, 94–101. http://doi.org/ https://doi.org/10.1016/j.livsci.2014.02.005.

Li, C., Zhang, G. F., Mao, X., Wang, J. Y., Duan, C. Y., Wang, Z. J., & Liu, L. B. (2016). Growth and acid production of Lactobacillus delbrueckii ssp. bulgaricus ATCC 11842 in the fermentation of algal carcass. Journal of Dairy Science, 99(6), 4243–4250. http://doi.org/10.3168/jds.2015-10700.

Li, J., Zhou, R., Ren, Z., Fan, Y., Hu, S., Zhuo, C., & Deng, Z. (2019). Improvement of protein quality and degradation of allergen in soybean meal fermented by Neurospora crassa. LWT, 101, 220–228. http://doi.org/ https://doi.org/10.1016/j.lwt.2018.10.089.

Schwendimann, L., Kauf, P., Fieseler, L., Gantenbein-Demarchi, C., & Miescher Schwenninger, S. (2015). Development of a quantitative PCR assay for rapid detection of Lactobacillus plantarum and Lactobacillus fermentum in cocoa bean fermentation. Journal of Microbiological Methods, 115, 94–99. http://doi.org/10.1016/j.mimet.2015.05.022.

Sheu, S., Hwang, W., Chiang, Y., Lin, W., Chen, H., & Tsen, H. (2010). Use of tuf gene-based primers for the PCR detection of probiotic Bifidobacterium species and enumeration of bifidobacteria in fermented milk by cultural and quantitative real-time PCR methods. Journal of Food Science, 75(8), M521–M527. http://doi.org/10.1111/j.1750-3841.2010.01816.x.

Shi, C., Zhang, Y., Lu, Z., & Wang, Y. (2017). Solid-state fermentation of corn-soybean meal mixed feed with Bacillus subtilis and Enterococcus faecium for degrading antinutritional factors and enhancing nutritional value. Journal of Animal Science and Biotechnology, 8(1), 50. http://doi.org/10.1186/s40104-017-0184-2.

Wang, Z., Li, Y., Jiang, L., Qi, B., Zhou, L., & Hernández-Ledesma, B. (2014). Relationship between Secondary Structure and Surface Hydrophobicity of Soybean Protein Isolate Subjected to Heat Treatment. Journal of Chemistry, 2014, 475389. http://doi.org/10.1155/2014/475389.

Yasar, S., Tosun, R., & Sonmez, Z. (2020). Fungal fermentation inducing improved nutritional qualities associated with altered secondary protein structure of soybean meal determined by FTIR spectroscopy. Measurement, 161, 107895. http://doi.org/10.1016/j.measurement.2020.107895.

Zhao, S., Li, P., Zhang, Q., Zhang, J., & Kong, L. (2013). Production of reducing sugar from corn stover by dilute acid hydrolysis co-catalyzed with metal salts under microwave radiation. Research On Chemical Intermediates, 59(8), 3803–3812. http://doi.org/10.1007/s11164-012-0882-5.

Zhou, Y., Zeng, Z., Xu, Y., Ying, J., Wang, B., Majeed, M.,… Li, W. (2020). Application of Bacillus coagulans in Animal Husbandry and Its Underlying Mechanisms. Animals (Basel), 10(3), 454. http://doi.org/10.3390/ani10030454.

